# Phthalate exposure influences mating behavior and sperm morphology in an aquatic ecotoxicology model system

**DOI:** 10.1101/2024.08.20.608834

**Authors:** Bryan Lamberto Guevara, Nadia Patel, Yi Tu, Maurine Neiman

## Abstract

Phthalates are a group of chemicals used to make plastics more durable, found in applications from cosmetics, lubricating oils, and flooring to soap, shampoo, and hairspray (CDC, 2021). Phthalates are also now known to be endocrine disruptors with connections to adverse reproductive outcomes in animals, including humans. Here, we evaluate the potential effects of a widely used phthalate ester, dimethyl phthalate (DMP), on male reproduction in a freshwater snail. DMP is found in industrial applications like solid rocket propellant as well as consumer products such as insect repellents and plastics. While there is some evidence that DMP negatively affects reproduction, especially in females, we still know very little about potential DMP effects on males. We addressed this important knowledge gap by testing the effects of DMP on *Potamopyrgus antipodarum*, a prosobranch snail native to New Zealand. These snails are very sensitive to water conditions and environmental chemicals, including endocrine-disrupting compounds, and are thus rising in prominence as water-quality sentinels and ecotoxicology models. We exposed experimental groups of male *P. antipodarum* to one of three different concentrations of DMP and characterized mating behavior and sperm morphology as a function of DMP exposure. Differences in these traits were primarily observed between the males in the control versus the High (10^-6^ M) DMP concentration group. As DMP exposure levels increased, we found that mating frequency ultimately decreased by more than 69% and that sperm morphology was increasingly altered relative to control males. Altogether, study outcomes suggest DMP exposure in male animals could have negative effects on reproduction, with particular relevance in aquatic and marine environments that are especially likely to harbor leached endocrine-disrupting chemicals.

## Introduction

Plastic waste is a major global issue, harming everything from marine organisms like turtles and seabirds to terrestrial organisms such as elephants and cattle. While plastics are capable of deteriorating, this process takes much longer than for organic matter. This extended process of decomposition leads to the formation of microplastics that can make their way into our rivers, streams, and waterways, ultimately endangering humans as well as aquatic and marine ecosystems (Ragusa et al. 2021). For example, we now know that microplastics can be found in all stages of human life, from prenatal development (Ragusa et al. 2021) to late adulthood.

In addition to posing physical harm, many microplastics also harbor endocrine-disrupting chemicals (EDCs) that are added to commercial plastics as plasticizers (D’Angelo & Meccariello 2021). There is emerging evidence that EDCs can damage human reproductive health as they accumulate in the body (Cuenca 2020). EDCs leach when they dissociate from the rest of the polymer as a consequence of environmental stressors such as temperature and pH changes (D’Angelo & Meccariello 2021). Consumer plastics are commonly exposed to UV radiation, humidity, and abrasion, which can exacerbate EDC leaching in marine environments (Chen et al. 2019).

Once leached, EDCs can enter human systems orally, topically, and via inhalation, and can then affect the endocrine system by, for example, mimicking natural hormones or blocking hormone access to intended receptors and signaling mechanisms (Diamanti-Kandarakis et al. 2009). In humans, EDCs have been linked to abnormalities in sex organs, certain cancers, metabolic issues, immune function, and decreased sperm quality and overall fertility in both males and females (Pan et al. 2024). Considered broadly, the growing environmental prevalence of EDCs has the capacity to alter sex ratios, impair immune function, disrupt reproductive health, and ultimately cause population declines (Windsor et al. 2018).

A class of EDCs known as phthalates are of particular concern as they are not chemically bound to the plastic polymer, increasing the likelihood of leaching when faced with environmental stress. Phthalates have been used since the 1930s as plasticizers for PVC plastics, medical instruments, children’s toys, and personal care products. Phthalates are also found in a variety of different commercial products, with the implication that phthalates are more likely to bioaccumulate and at a greater rate than other common commercially used EDCs (Heudorf et al. 2007). The negligent production, use, and disposal practices of plastic-based products are the major source of phthalates in aquatic systems via runoff and leaching (Metcalfe et al. 2022). While we still know relatively little about consequences of phthalates for humans, increased phthalate exposure has been associated with decreased testosterone levels and increased levels of estradiol in male humans, which could in turn reduce male fertility (Zhu et al. 2022).

Dimethyl phthalate (DMP) is a phthalate ester (PAE) found in various consumer plastics. Most notably, DMP is a main component of plastic #1, which is primarily polyethylene terephthalate (PET). DMP, despite its similarities to other more toxic PAEs, has not been found to be toxic during development (Field et al. 1993, NTP 1989). We do know from animals experiencing experimental oral exposure to DMP that this exposure can translate into relatively minor decreases in kidney function (U.S. EPA 1987, U.S. DHHS 1993). What remains largely uncharacterized is whether environmental DMP influences reproduction. The few studies of which we are aware have produced mixed results. For example, while Gray et al. (2000) found that perinatal exposure to DMP did not affect testis weight, areola size, or frequency of reproductive malformations in mice, Mei et al. (2019) found that prolonged exposure to DMP for female mice decreased the secretion of follicle-stimulating hormones and increased the levels of the estradiol and luteinizing hormones that are critical regulators of the menstrual cycle in mammals. There is an especially obvious knowledge gap regarding whether and how DMP alters male reproductive traits, and especially after the postnatal period.

### Using a freshwater snail ecotoxicology model to test DMP effects on sperm

*Potamopyrgus antipodarum* is a New Zealand freshwater snail that inhabits a wide range of freshwater and brackish habitats and is commonly located in more disturbed areas (Loo et al. 2007, Ponder 1988, Zaranko et al. 1997). This snail is also a global invader (reviewed in Geist et al. 2024). The widespread distribution, broad habitat use, availability of asexual lineages (e.g., Möller et al. 1996), and documented sensitivity to environmentally relevant concentrations of EDCs (Duft et al. 2003, 2007, Jobling et al. 2003, Stange & Oehlmann 2012, Lecomte et al. 2013) make *P. antipodarum* a powerful ecotoxicology model and aquatic sentinel (e.g., Geiß et al. 2017, Subba et al. 2021). Most of the studies demonstrating sensitivity to EDCs and other toxicants in *P. antipodarum* emphasize their effects on female reproductive output or fecundity. By contrast, we know little about the potential for EDCs (including DMP) to affect reproductive function in male *P. antipodarum*.

A previous study in mice found evidence that motility and activity in sperm experiencing relatively short-term exposure to phthalates was reduced, a consequence of decreased sperm capacity to generate ATP when exposed to phthalates. This short-term phthalate exposure also resulted in abnormal capacitation, the penultimate step in maturation of mammalian spermatozoa (Amjad et al. 2021). A study of marine mussels exposed to dibutyl phthalate (DBP), di(2-ethylhexyl) phthalate (DEHP), and DMP reported increased expression of glutathione (GSH) and other antioxidant enzymes that protect cells against reactive oxygen species, relative to unexposed individuals. GSH acted as the primary antioxidant in the mussels exposed to DMP. This influx of GSH ultimately disturbed the metabolomic balance in these mussels, providing a mechanistic basis for a connection between DMP exposure and sperm function (Gu et al. 2021). These previous studies set the stage to investigate whether and how EDCs affect male *P. antipodarum*, which can in turn provide insights into how these common environmental pollutants affect male function across taxa.

In accordance with Dalsenter et al. (2006), who found increases in intromission latency and number of intromissions for ejaculation for male mice with exposure of 500 mg/kg per day of another phthalate, we hypothesize that DMP will depress male *P. antipodarum* mating behavior. Using a rat model, Parks et al. (2000) and Drake et al. (2009) saw decreased fetal testosterone synthesis and disruption of testosterone production in male reproductive development when the rats were exposed to DEHP and DBP, respectively, leading us to hypothesize that phthalate exposure will alter sperm production in *P. antipodarum* via its consequences for testosterone signaling. We evaluated these hypotheses by comparing the mating behavior and sperm morphology of male *P. antipodarum* that we experimentally exposed to one of three different concentrations of DMP, predicting that mating frequency and duration will decrease and sperm morphology will become increasingly deviant relative to controls as DMP concentration increases.

## Methods

### Snail selection

*Potamopyrgus antipodarum* males used in this study had been cultured under laboratory conditions for several years at the University of Iowa following descent from a diverse set of sexual snails collected from multiple New Zealand lake populations. These snail cultures are housed in 20-liter tanks filled with carbon-filtered tap water in a 16° C room with a 16:8-hr light:dark cycle and a relative humidity of 64%. Snails were haphazardly selected from these tanks. Males were then distinguished from females by visualizing each snail under a NIKONSMZ800 stereo microscope with a magnification ranging from 10x to 63x and then by turning each individual on its back and waiting for the snail to right itself, which reveals the penis on the right side of the male body. We also measured the length of each snail at this time, choosing only snails that were =\> 3.0 mm from apex to aperture, typically used as the lower size threshold for adulthood in this species (e.g., Larkin et al. 2016).

### Experimental Design

Each of the 32 males that we used for this experiment was placed individually into a 16 oz. glass Mason jar filled with 300 mL of carbon-filtered tap water. While we did not monitor water quality aside from regular quantification of chlorine levels, all water for all treatments came from the same source at the same time, allowing us to exclude water quality differences across treatments other than our focal DMP treatments as the source of any treatment effects. We applied one of three DMP treatments to each of 24 snails (8 snails per treatment), with the remaining 8 snails serving as controls (no DMP). The DMP-exposed snails were housed in water containing 10^-10^ M DMP (“low” treatment), 10^-8^ M DMP (“intermediate” treatment), or 10^-6^ M DMP (“high” treatment). The snails in the control group did not receive any experimental DMP exposure. Because we were not aware of any other studies that manipulated DMP exposure in snails, we based our DMP exposure levels on concentrations used for another common household EDC, bisphenol A, in a study of its effects on embryo production in *P. antipodarum* (Jobling et al. 2003). We broadened the range of concentrations employed by Jobling et al (2003) in order to increase our likelihood of detecting treatment effects. The snails were exposed to these DMP concentrations for a five-week period prior to mating trials and dissection for sperm characterization. We used this timeline because spermatogenesis generally takes 2 – 5 weeks under optimal conditions in small gastropods (de Freitas Tallarico et al. 2004, Rodrigues et al. 2021). Following standard *P. antipodarum* culture conditions (e.g., Stork et al. 2022), each jar of snails was supplied 260 µL of a solution containing 50 mL carbon-filtered tap water, 0.096 g of freshwater fish fry food, and 0.01 g of chalk 3x/week.

### Mating Trials

Following the procedures for mating trials used in a previous study of *P. antipodarum* mating behavior (Stork et al. 2022), an individual trial consisted of one male and two female *P. antipodarum* placed into a 100-mm petri dish. Male snails were marked with nail polish on their shells to readily differentiate them from females. The shallow structure and relatively small size of the petri dish has been found to be an effective means of ensuring that snails come in contact during the two hours of the trial along with facilitating clear video analysis of behavior (Stork et al. 2022). We used a two-hour time period for the trial following Neiman & Lively (2005), who showed that *P. antipodarum* copulation is almost always concluded within minutes to an hour. At the start of the two-hour period, the two females were placed equidistant from the male. Next, all three snails were allowed to freely move around the dish for two hours. We filmed the petri dish from above using a 1080p Logitech Webcam and the time-lapse software VideoVelocity (v3.7.2104; CandyLabs 2022). Following Nelson & Neiman (2011) and Stork et al. (2022), we defined mating attempts as a physical interaction involving one snail mounting another for more than 15 seconds (Nelson & Neiman 2011, Stork et al. 2022). We used these recordings to quantify the frequency of mating attempts per male and the duration of each attempt.

### Dissections & Microscopic Visualization

We first dissected each male by using fine-tipped forceps to break the shell at its widest point, enabling careful extraction of the whole body and leaving the sperm duct intact. We then identified and removed the sperm duct, wet mounted the sperm duct on a microscope slide, and sealed the slide with acrylic paint to retain moisture. We visualized sperm under a Leica DMi8 automated microscope using brightfield technology. Following a similar study of *P. antipodarum* sperm by Jalinsky et al. (2020), we took images of ∼10 sperm per snail at 1000x and used these images and ImageJ software (Schneider et al. 2012) to measure head length, width, circumference, and sperm tail length (Figure 1).

**Figure 1.**
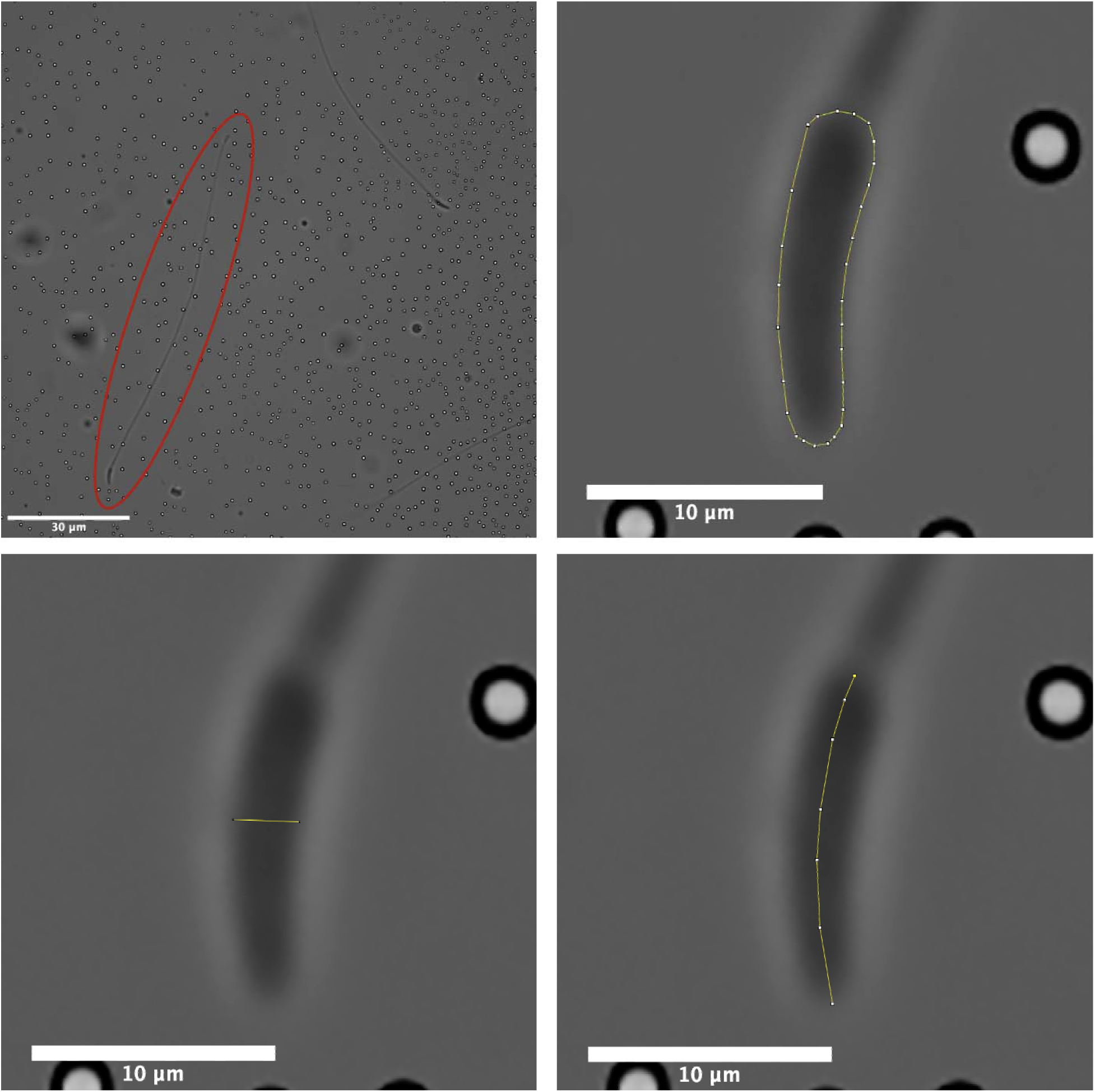
Sperm structure and microscopic imaging and measurement in *P. antipodarum*. Black bar scaled to 30 µm. Yellow segmented line represents the measuring tool used for measuring sperm parameters. Top left panel depicts isolated sperm at 10000x with red circle surrounding sperm selected for measurement. Top right, bottom left, and bottom right panel depict how sperm head circumference, width, and length, respectively, were measured in ImageJ.

### Statistical Analyses

We used Shapiro-Wilks tests to assess normality for each dataset (mating frequency, mating duration, sperm head length, sperm head width, sperm head circumference, sperm head length, sperm tail length), and found that each of these datasets represented significant departures from normality (*p* < 1.01e^-05^). Accordingly, we used nonparametric approaches for all subsequent analyses: Kruskal-Wallis tests for the comparisons of mating attempt frequency, mating attempt duration, and sperm head length, width, and circumference, and tail length across the four DMP treatment groups. We also compared each of the possible pairwise sets from the Kruskal-Wallis test using the post-hoc Dunn test. All data analyses were performed with R (R Core Team, 2024) via janitor (v.2.2.0; Firke 2023), dplyr (v1.1.4; Wickham et al. 2023), and FSA (v0.9.5; Ogle et al. 2023). Data were visualized with ggplot2 (Wickham 2016) and ggstatsplot (Patil 2021).

## Results

The mating behavioral trials revealed that mating frequency differed across the treatment groups (*X^2^* = 15.4, df = 3, *p* = 0.001), with mating frequency in the control group significantly higher than both the intermediate and high DMP treatment groups (Figure 2, Supplementary Figure 1, Supplementary Table 1). We did not observe any significant differences between treatment groups for mating duration (*X^2^* = 5.62, df = 3, *p* = 0.132; Supplementary Table 1).

**Figure 2.**
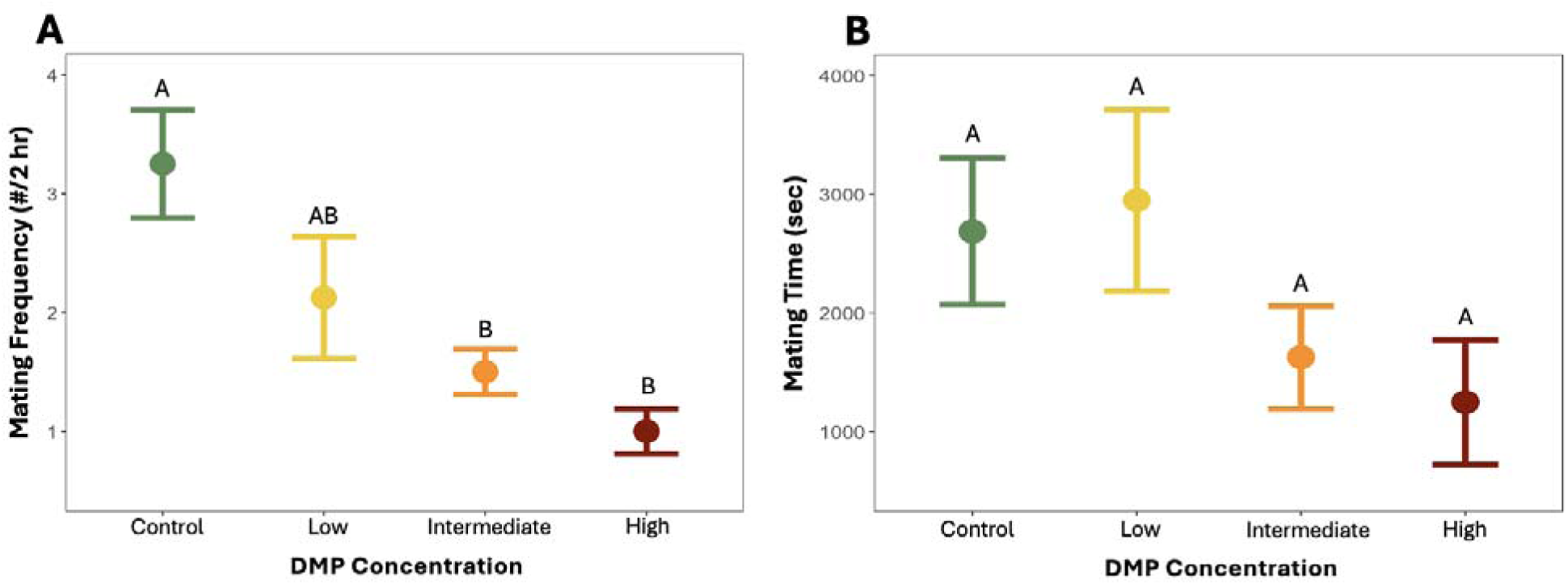
Increasing DMP concentration is associated with decreased mating frequency and duration in male *P. antipodarum*. Comparisons of mean (A) frequency of male mating across the 4 DMP treatment groups and (B) time males spent mating. Error bars represent standard error. Statistically significant different pairwise comparisons are denoted by different versus shared letters above error bars.

There were no significant differences across the four DMP treatment groups for sperm head circumference and head length (Table 1, Figure 3, Supplementary Figure 2, Supplementary Table 2). The treatment groups did differ in sperm head width and tail length (Supplementary Table 2). Post-hoc pairwise analysis revealed that this outcome was driven by the significantly shorter sperm head width in the high DMP group vs. each of the other 3 DMP treatments. There were no significant differences between the intermediate and low, intermediate and control, and control and low DMP treatment groups for sperm head width. Snails from the control treatment had significantly longer tails than snails from the intermediate and high DMP treatments. Additionally, snails from the low DMP treatment had longer tails than snails exposed to high DMP. These results are summarized in Supplementary Table 3.

**Figure 3.**
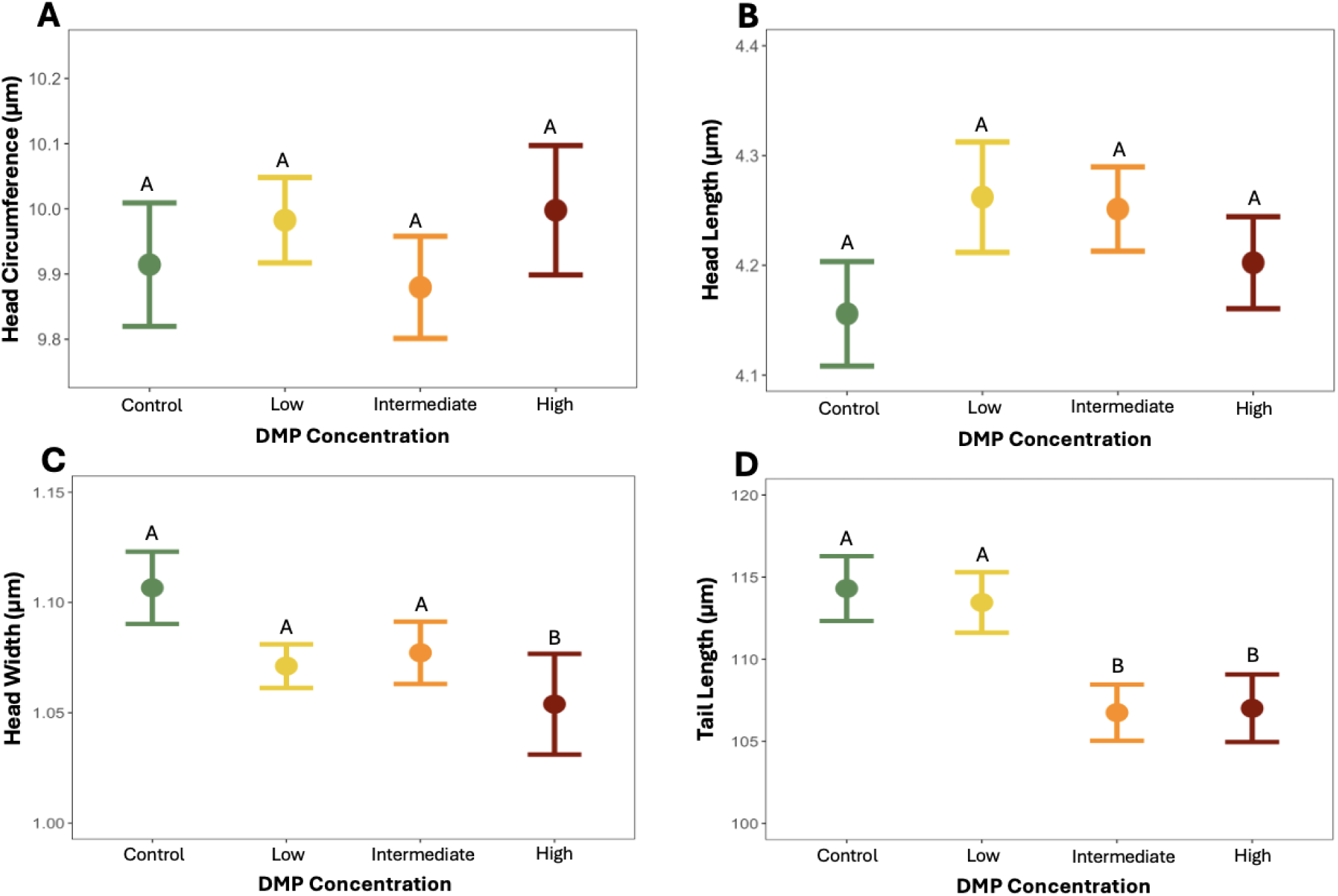
Sperm morphology as a function of DMP concentration. Comparisons of mean sperm (A) head circumference, (B) head length, (C) head width, and (D) tail length across varying levels of DMP. Error bars represent standard error. Statistically significant different pairwise comparisons are denoted by different versus shared letters above error bars.

**Table 1.** Mean and standard deviation (SD) of sperm metrics from 32 *P. antipodarum* males across the four treatment groups in micrometers (µm).

| DMP Treatment | Head Circumference (SD) | Head Length (SD) | Head Width (SD) | Tail Length (SD) |
| --- | --- | --- | --- | --- |
| Control | 9.91 (0.79) | 4.16 (0.40) | 1.11 (0.14) | 114.31 (16.45) |
| Low | 9.98 (0.59) | 4.26 (0.45) | 1.07 (0.09) | 113.46 (16.49) |
| Intermediate | 9.88 (0.66) | 4.25 (0.32) | 1.08 (0.12) | 106.75 (14.30) |
| High | 10 (0.83) | 4.20 (0.35) | 1.06 (0.19) | 107.01 (17.19) |

## Discussion

We used an increasingly prominent ecotoxicology model system, *Potamopyrgus antipodarum*, to evaluate potential consequences for male reproduction of exposure to a common household EDC, dimethyl phthalate (DMP). Our study is novel in comparing mating behavior and sperm morphology across various levels of DMP concentration in a freshwater mollusk, which allows us to observe the short-term effects that exposure to EDCs may impose on male reproduction.

Our experiment revealed that increased exposure to DMP resulted in a significant decrease in male mating frequency and a trend towards a reduction in mating duration. Increased DMP concentration was also associated with significantly shorter sperm tails and head widths. While the relationship between sperm morphology and fertility is a contentious topic, there is evidence from multiple animal systems that do demonstrate that sperm morphology is associated with reproductive hormone levels (Zhao et al. 2020) and the extent of fragmentation of sperm nuclear DNA (Gu et al. 2021), which are in turn linked to fertility (Benchaib et al. 2003). To give a specific example relevant to our own work, Oppliger et al. (2003) and Radwan (1996) showed that sperm tail length is a critical predictor of motility and ability to successfully fertilize. Alterations to the sperm head shape or size may be indicative of DNA damage, as revealed by studies indicating that expression of GSH, an antioxidant that protects cells against damage from reactive oxygen species, is increased when exposed to phthalates (Gu et al. 2021). Studies in biological models such as the pig also demonstrate that sperm morphology affects sperm motility characteristics (Gil et al. 2009).

Our results are consistent with a scenario where DMP exposure could indeed influence male fertility in freshwater organisms like *P. antipodarum* and perhaps in other, more distantly related animals like humans. Because, to the best of our knowledge, no studies have gone beyond showing increased frequency of morphological abnormalities in sperm produced by males exposed to EDCs (Hauser et al. 2003, Pérez-Herrera et al. 2008, Dziewirska et al. 2018), whether these morphological and behavioral changes we observed will influence can affect male fertility remains to be seen. Fruitful next steps could include assessing the expression and release of antioxidants in *P. antipodarum* when exposed to phthalates or other EDCs, which may reveal DNA damage and provide insight regarding the ability of damaged sperm ability to fertilize. Direct observations of motility as a function of changes in *P. antipodarum* sperm tail length would provide further insight into potential links between phthalate exposure and fertility.

These effects of DMP exposure were evident when DMP treatments were confined to *adult* snails is notable in light of the fact that phthalate exposure early in development can cause testicular dysgenesis syndrome (TDS) in humans (Hlisníková et al. 2020). With this point in mind, it would be valuable to experimentally expose juvenile *P. antipodarum* males to phthalates to determine whether gonadal structure and function are affected and whether and how these consequences of exposure affect sperm morphology and function. To date, and to the best of our knowledge, the potential for contaminant-induced abnormalities in mature sperm has not been studied in any marine or aquatic invertebrate species.

It is possible that at least some of the mating and sperm-related consequences of DMP exposure for adult snails might be a function of stress. Indeed, chronic stress is known to lower testosterone levels and damage reproductive organs in humans (Xiong et al. 2022). In humans, lifestyle factors such as obesity, prolonged exposure to mobile phone radiation, and occupational stress have been shown to significantly damage sperm DNA and result in immature sperm (Radwan et al. 2016). Mental and physical stress are also known to enhance the presence of free radicals, ultimately leading to oxidative stress that can in turn result in reduced sperm count and sperm motility in humans (Nowicka-Bauer & Nixon 2020). Thermal stress also decreases mating frequency and inhibits flight in *Drosophila mojavensis,* but not courtship behaviors such as wing waving (Patton & Krebs 2011). At least some of these phenomena linked to thermal stress could be linked to the activation of heat shock protein (HSP; and specifically HSP70). Indeed, high levels of HSP70 have been reported in infertile men relative to fertile men (Erata et al. 2008, Kumar & Singh 2022), with Erata et al. (2008) concluding that HSP70 expression increased in these infertile men as a defense mechanism against sperm apoptosis.

Together, our study suggests that DMP exposure for adult male animals can manifest in altered mating behavior and sperm morphology. These results are significant in being documented in an ecotoxicology and aquatic sentinel model system, suggesting that the endocrine-disrupting chemicals that leach from plastics into aquatic and marine systems may be disrupting male fertility in the organisms that inhabit these environments.

### Implications for Conservation

We provide new insight into the potential threat that plastic pollution poses for the reproductive health of aquatic organisms. We revealed that exposure to a common EDC, generally assumed to be relatively harmless for humans, alters mating behavior and sperm morphology in freshwater snails. Because *P. antipodarum* serves as an aquatic sentinel and ecotoxicology model system, our results suggest that plastic pollution, through the leaching of EDCs into water bodies, could disrupt reproduction of various aquatic organisms. Over extended periods of time, the leaching of these chemicals into our water bodies may lead to population decline for organisms living in exposed habitats, with potential cascading effects on biodiversity and ecosystem services.

These findings highlight the need for conservation efforts to prioritize plastic pollution control and monitoring for EDCs to protect the reproductive health of aquatic organisms and to ensure long term sustainability of the ecosystems they inhabit.

## Ethics and Integrity Statements

### Data Availability Statement

We will make all datasets publicly available upon publication in Dryad.

### Ethics and permit approval statement

We have the appropriate permit from the Iowa Department of Natural Resources to house and culture *P. antipodarum* in the Neiman lab.

### Permission to reproduce material from other sources

Not applicable.

## Acknowledgments

We gratefully acknowledge help and generosity with respect to microscopy from Dr. Sarit Smolikove and her lab group, funding from Linda and Rick Maxson, the Belin-Blank Center, the National Science Foundation UI-LSAMP program, and the Iowa Sciences Academy, feedback and input from the Neiman lab, and snail care from Morgan Anderson. We are also grateful for constructive feedback from several anonymous peer reviewers.

