## Supplementary Figure 1 for "Phthalate exposure influences mating behavior and sperm morphology in an aquatic ecotoxicology model system"

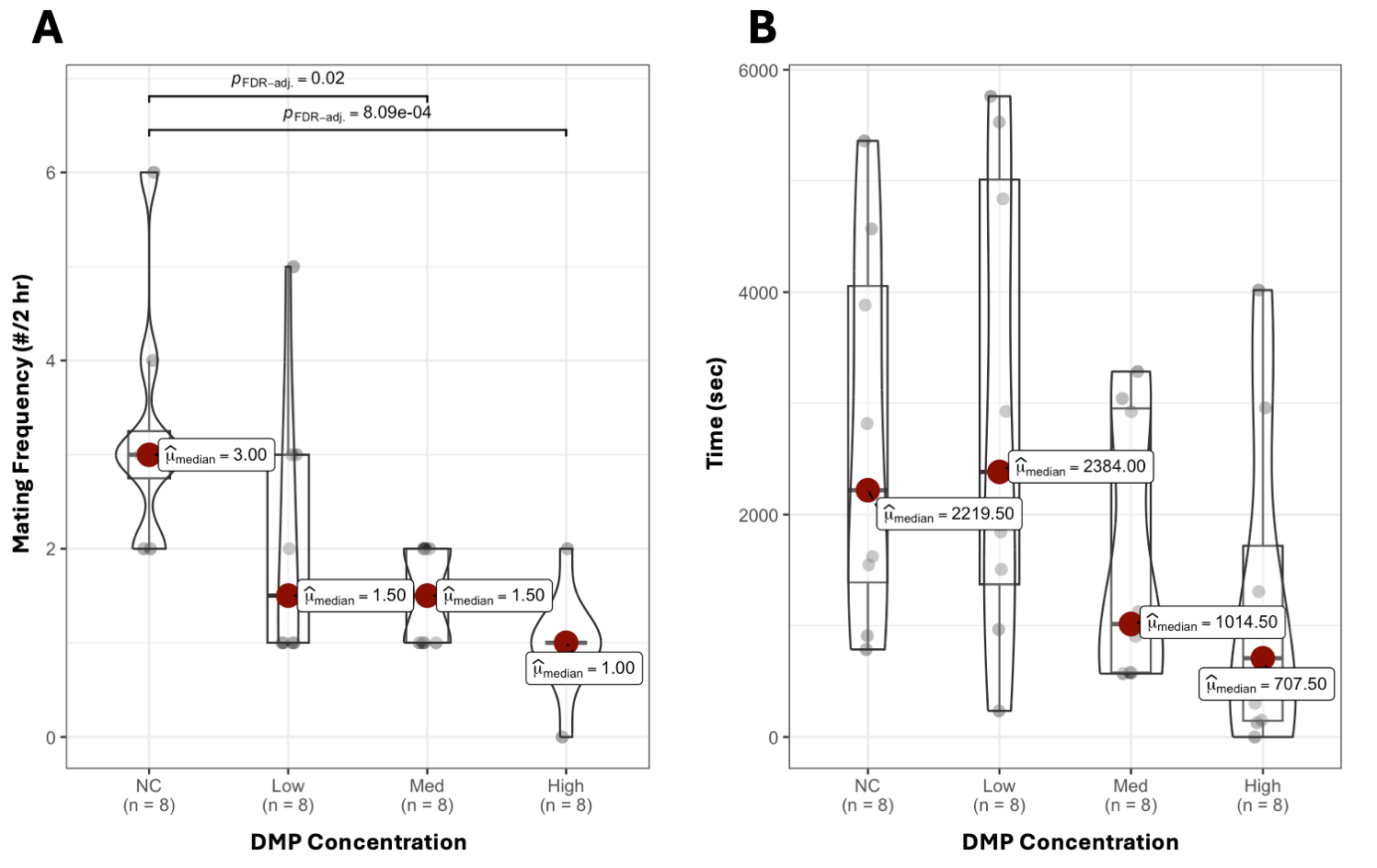

**Supplementary Figure 1**. Violin plots comparing (A) how frequently the males mated across the 4 DMP treatment groups and (B) time that males spent mating. Individual data points are indicated with black dots. Red dots highlight median and median values per treatment group. Statistically significant different pairwise comparisons are denoted above the groups compared with rounded FDR-adjusted *p*-values.

with black dots. Red dots highlight median values per treatment group. Statistically significant differences between pairwise comparisons are denoted above the groups.
