## Supplementary Figure 2 for "Phthalate exposure influences mating behavior and sperm morphology in an aquatic ecotoxicology model system"

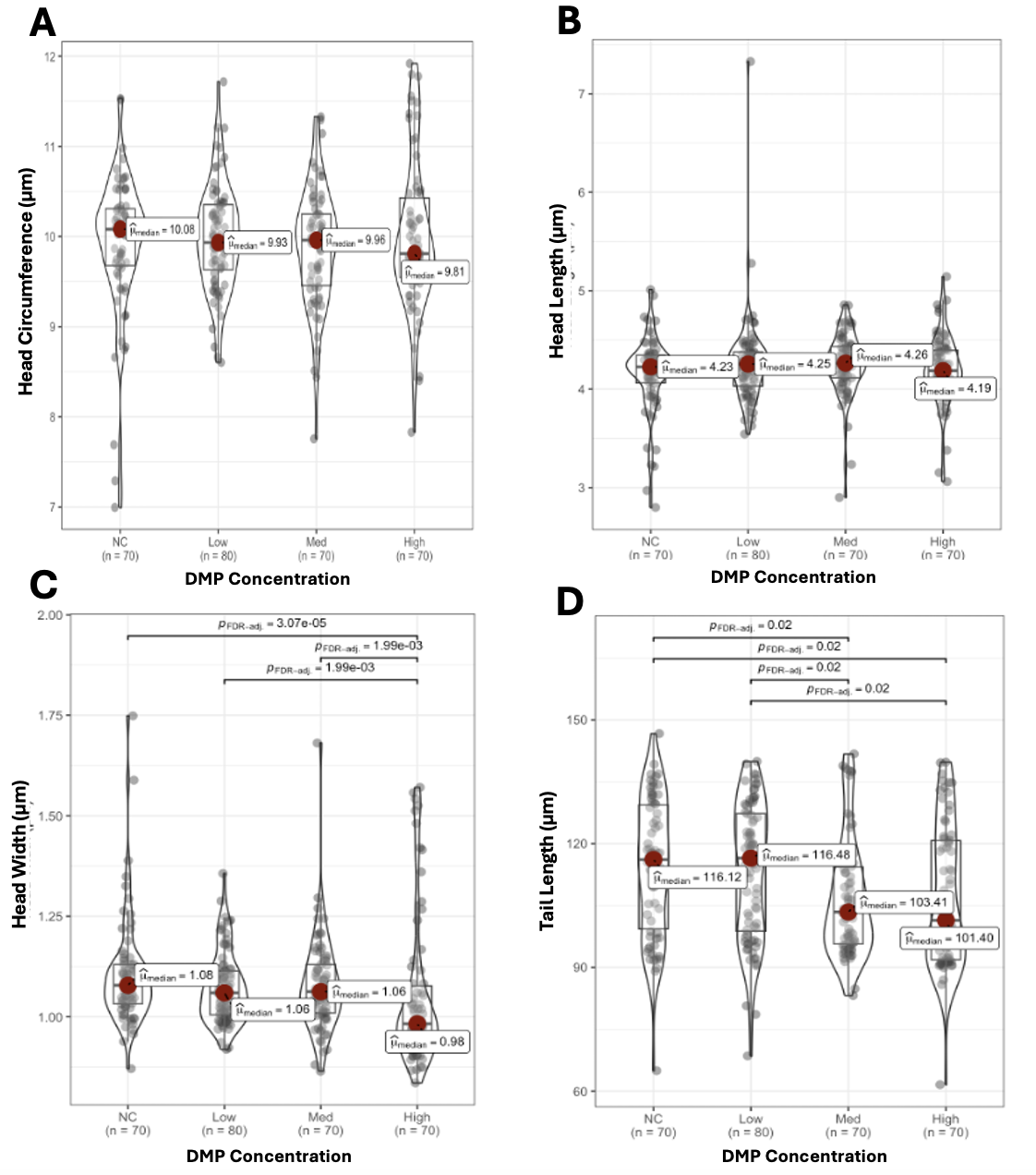

**Supplementary Figure 2. Sperm morphology as a function of DMP concentration**. Violin plots comparing sperm (A) head circumference, (B) head length, (C) head width, and (D) tail length across varying levels of DMP. Individual data points are indicated with black dots. Red dots highlight median values per treatment group. Statistically significant differences between pairwise comparisons are denoted above the groups.
