## Supplementary Table 1 for "Phthalate exposure influences mating behavior and sperm morphology in an aquatic ecotoxicology model system"

Supplementary table 1. Dunn pairwise posthoc analysis for mating frequency. Significant results are **bolded.**

|  | Treatment comparisons | z | *p* |
| --- | --- | --- | --- |
| Mating frequency | Control vs. Low | -1.96 | **0.001** |
|  | Control vs. Intermediate | -2.68 | **0.022** |
|  | Control vs. High | -3.82 | **<0.001** |
|  | Low vs. Intermediate | 0.718 | 0.472 |
|  | Low vs. High | -1.86 | 0.094 |
|  | Intermediate vs. High | -1.14 | 0.305 |
