## Supplementary Table 2 for "Phthalate exposure influences mating behavior and sperm morphology in an aquatic ecotoxicology model system"

Supplementary table 2. Kruskal-Wallis analysis statistics for sperm morphology. Significant results are **bolded**.

| Sperm morphology | *X^2^* | df | *p* |
| --- | --- | --- | --- |
| Head circumference | 1.23 | 3 | 0.744 |
| Head length | 2.56 | 3 | 0.464 |
| Head width | 22.9 | 3 | **<0.001** |
| Tail length | 14.6 | 3 | **0.002** |
