## Supplementary Table 3 for "Phthalate exposure influences mating behavior and sperm morphology in an aquatic ecotoxicology model system"

Supplementary table 3. Dunn-test pairwise analysis for sperm head width and tail length. Significant results are **bolded**.

| Sperm morphology | Treatment comparisons | z | *p* |
| --- | --- | --- | --- |
| Head width | Control vs. Low | 1.42 | 0.234 |
|  | Control vs. Intermediate | -1.22 | 0.266 |
|  | Control vs. High | -4.55 | **<0.001** |
|  | Low vs. Intermediate | -0.156 | 0.875 |
|  | Low vs. High | -3.29 | **0.002** |
|  | Intermediate vs. High | -3.34 | **0.003** |
| Tail length | Control vs. Low | -0.163 | 0.870 |
|  | Control vs. Intermediate | -2.64 | **0.016** |
|  | Control vs. High | -2.82 | **0.028** |
|  | Low vs. Intermediate | 2.57 | **0.018** |
|  | Low vs. High | -2.75 | **0.009** |
|  | Intermediate vs. High | -0.177 | 1.00 |
